# Seasonal patterns of dengue fever in rural Ecuador: 2009—2016 Seasonality of dengue fever in Ecuador

**DOI:** 10.1101/452318

**Authors:** Rachel Sippy, Diego Herrera, David Gaus, Ronald E. Gangnon, Jonathan A. Patz, Jorge E. Osorio

## Abstract

Season is a major determinant of infectious disease rates, including arboviruses spread by mosquitoes, such as dengue, chikungunya, and Zika. Seasonal patterns of disease are driven by a combination of climatic or environmental factors, such as temperature or rainfall, and human behavioral time trends, such as school year schedules, holidays, and weekday-weekend patterns. These factors affect both disease rates and healthcare-seeking behavior. Seasonality of dengue fever has been studied in the context of climatic factors, but short- and long-term time trends are less well-understood. With 2009—2016 medical record data from patients diagnosed with dengue fever at two hospitals in rural Ecuador, we used Poisson generalized linear modeling to determine short- and long-term seasonal patterns of dengue fever, as well as the effect of day of the week and public holidays. In a subset analysis, we determined the impact of school schedules on school-aged children. With a separate model, we examined the effect of climate on diagnosis patterns. In the first model, the most important predictors of dengue fever were annual sinusoidal fluctuations in disease, long-term trends (as represented by a spline for the full study duration), day of the week, and hospital. Seasonal trends showed single peaks in case diagnoses, during mid-March. Compared to the average of all days, cases were more likely to be diagnosed on Tuesdays (risk ratio (RR): 1.26, 95% confidence interval (CI) 1.05—1.51) and Thursdays (RR: 1.25, 95% CI 1.02—1.53), and less likely to be diagnosed on Saturdays (RR: 0.81, 95% CI 0.65—1.01) and Sundays (RR: 0.74, 95% CI 0.58—0.95). Public holidays were not significant predictors of dengue fever diagnoses, except for an increase in diagnoses on the day after Christmas (RR: 2.77, 95% CI 1.46—5.24). School schedules did not impact dengue diagnoses in school-aged children. In the climate model, important climate variables included the monthly total precipitation, an interaction between total precipitation and monthly absolute minimum temperature, an interaction between total precipitation and monthly precipitation days, and a three-way interaction between minimum temperature, total precipitation, and precipitation days. This is the first report of long-term dengue fever seasonality in Ecuador, one of few reports from rural patients, and one of very few studies utilizing daily disease reports. These results can inform local disease prevention efforts, public health planning, as well as global and regional models of dengue fever trends.

**Author summary:** Dengue fever exhibits a seasonal pattern in many parts of the world, much of which has been attributed to climate and weather. However, additional factors may contribute to dengue seasonality. With 2009— 2016 medical record data from rural Ecuador, we studied the short- and long-term seasonal patterns of dengue fever, as well as the effect of school schedules and public holidays. We also examined the effect of climate on dengue. We found that dengue diagnoses peak once per year in mid-March, but that diagnoses are also affected by day of the week. Dengue was also impacted by regional climate and complex interactions between local weather variables. This is the first report of long-term dengue fever seasonality in Ecuador, one of few reports from rural patients, and one of very few studies utilizing daily disease reports. This is the first report on the impacts of school schedules, holidays, and weekday-weekend patterns on dengue diagnoses. These results suggest a potential impact of human behaviors on dengue exposure risk. More broadly, these results can inform local disease prevention efforts and public health planning, as well as global and regional models of dengue fever trends.

## Introduction

Seasonality of infectious disease is a phenomenon commonly observed in the northern and southern hemispheres, with seasonality of influenza being the most well-known and well-studied infectious disease with a seasonal pattern [1-6]. Seasonality has also been observed with other infectious diseases, including malaria [7], dengue [8], tuberculosis [9, 10], acute respiratory infection [1, 11], and foodborne illness [12-15]. These relationships are often a combination of climatic and environmental factors and how these factors affect pathogen transmissibility [15, 16], vector abundance [8, 17-21], and human health, and drive human behaviors such as diet, crowding, travel patterns, and outdoor exposures [8, 14, 15, 19, 20].

Mosquito-borne viral infections include dengue fever, yellow fever, chikungunya, and Zika, among others [22]. These illnesses are common in tropical countries and are most often spread by mosquitoes in the *Aedes* genus. Dengue virus is the most common, and may present with fever, rash, and general pain; although an estimated 80% of dengue patients are asymptomatic [23], this infection can have serious health consequences, including death [24].

The diagnosis of dengue and other acute febrile illnesses can be extremely difficult, depending on the stage of the illness and the resources available at the point of care. Dengue cannot always be distinguished from other febrile illnesses, though diagnostic testing, including rapid tests, ELISA, and PCR-based assays are sometimes available and can aid with diagnosis [25], though the sensitivity and specificity of these tests are not perfect. Correct diagnosis of dengue additionally relies on the patient’s presenting signs and symptoms as well as the expertise of the clinician.

Seasonality affects dengue diagnosis rates through several mechanisms. Seasons drive human behavior: people may be more or less likely to spend time crowded indoors or spread outdoors depending on the time of year, which affects exposure rates. This can be the result of weather conditions or a result of seasonal holidays, which affect school and work schedules, and drive public gatherings (such as parades) or private family gatherings. There is also reason to believe that seasonality affects host immunity: in tropical countries, both cell-mediated and humoral immune responses are decreased during the rainy season [26]. This could be driven by seasonal variation in gene expression [27], levels of immune-modulators and blood cell composition [28], food availability, daylight exposure, and/or environmental exposures [26], though the causal direction of changes in the immune system, season, and seasonal disease is unclear. In addition, long-term or multi-annual disease trends are often a reflection of a buildup of disease-specific immunity in a population: for outbreaks to occur, there must be a sufficient number of susceptible individuals in the population. If all persons in the community were infected in the previous years and are therefore immune to circulating strains of virus, no outbreak occurs and the season will have a relatively low intensity, and the low intensity will continue until additional susceptibles are available from birth, migration, or introduction of a new dengue serotype.

Climate is a major component of seasonality and directly impacts the life history and behavior of the mosquito vector. *Aedes aegypt*i, which is the principal vector of dengue in Ecuador, has been well-characterized in its relationship to temperature, which has been shown to impact development rates, lifespan, fecundity, survival, biting rates, transmission probability, infection probability, abundance and incubation rates in both field and laboratory studies [29-36]. Field studies of rainfall have found associations between larval or adult abundance and precipitation [37-39]. Because temperature and precipitation can affect mosquitoes throughout their life course, the temporal scale of climate-mosquito associations can vary, depending on the life stage of the mosquito. For example, lagged precipitation (one to two months prior) is linked to larval indices due to the impact of precipitation on larval breeding sites [37], while both lagged temperature (4 weeks) and unlagged [*i.e*. current] mean temperature have been associated with adult abundance [39, 40]. Adult abundance and biting patterns are critical to dengue risk; climate plays a major role in the activity levels of these vectors [33].

The climate of Ecuador is highly diverse; though small in area, it contains 11 different Köppen-Geiger climate classifications, with the coast being generally classified as hot and semi-arid or tropical savanna climates, the central Andean range as oceanic or warm-summer Mediterranean climates, and the eastern rainforest as tropical rainforest climates [41]. Ecuador is also impacted by the El Niño/Southern Oscillation (ENSO) phenomenon in which the surface temperature of the Pacific ocean leads to periodic changes in regional weather patterns [42]. Specifically, an El Niño year will be warmer and wetter than average in Ecuador, and a La Niña year will be drier and cooler than average [42].

Studies of disease seasonality in tropical regions are limited. For mosquito-borne disease, previous research has largely focused on climatic and environmental variables, which directly affect vector abundance. In Ecuador, this research has been limited to two studies of dengue cases in coastal regions; In one study, minimum weekly temperature and mean weekly precipitation were shown to be strongly linked to weekly number of dengue cases [19]. A second study in the same area found that minimum weekly temperature, precipitation, and El Niño events were positively associated with dengue risk [20]. These studies both occurred in a large city the southern coast of Ecuador; given the diversity of climates and communities in Ecuador and the need for relevant evidence to make policy decisions, it is important to determine if the causal relationships between seasonal factors, climates, and dengue cases are similar in other areas of Ecuador.

With the present study we determined the seasonality of dengue fever by decomposing seasonality into two components: non-climate seasonality and climate-driven seasonality, using data from patients clinically diagnosed with dengue fever at two hospitals in rural Ecuador with a subtropical climate. Non-climate trends included short- and long-term trends, and the effects of school sessions, public holidays, and weekdays on these diagnoses. Climate-driven trends included an examination of regional and local climate variable impacts on dengue fever diagnoses.

## Methods

### Study population & site

Hospital Pedro Vicente Maldonado (HPVM) is a 17-bed rural hospital located in Pedro Vicente Maldonado (PVM), Pichincha, Ecuador (Fig 1). It primarily serves patients from Cantons Pedro Vicente Maldonado, Puerto Quito, San Miguel de los Bancos, and Santo Domingo. Pedro Vicente Maldonado is located at 0°05′12.3″N, 79°03′08.0″W, and northwest of Quito, at approximately 600 meters altitude, with a projected 2016 population of 6,944. Hospital Saludesa (HS) is a 60-bed metropolitan hospital located in Santo Domingo de los Tsáchilas (SD), Santo Domingo de los Tsáchilas, Ecuador (Fig 1). It serves patients from Santo Domingo de los Tsáchilas Province. Santo Domingo de los Tsáchilas is located at 0°15′15″S, 79°10′19″W, and west of Quito, at approximately 550 meters altitude, with a population of 305,632 (2010 Census). Both hospitals have 24-hour, 7-days-a-week emergency rooms, with regular consultation available on Mondays—Saturdays. During holidays, only the emergency room services are available. Both hospitals have clinical laboratory services available, including the NS1 dengue antigen rapid and dengue IgG antibody rapid tests (Human, Wiesbaden, Germany); the NS1 dengue antigen rapid test is the diagnostic of choice. These cities have a tropical rainforest climate; average monthly temperatures run from 71.8° Fahrenheit (22.1° Celsius) in November to 74.8° Fahrenheit (23.8° Celsius) in March. Average total monthly precipitation runs from 110 millimeters (mm) in July to 671 mm in April. Both sites have ongoing mosquito control programs. Cities are fumigated approximately once per month with repellent, and residents are provided with temephos (Abate®) treatment for water stored in large laundry tanks.

**Fig 1.**
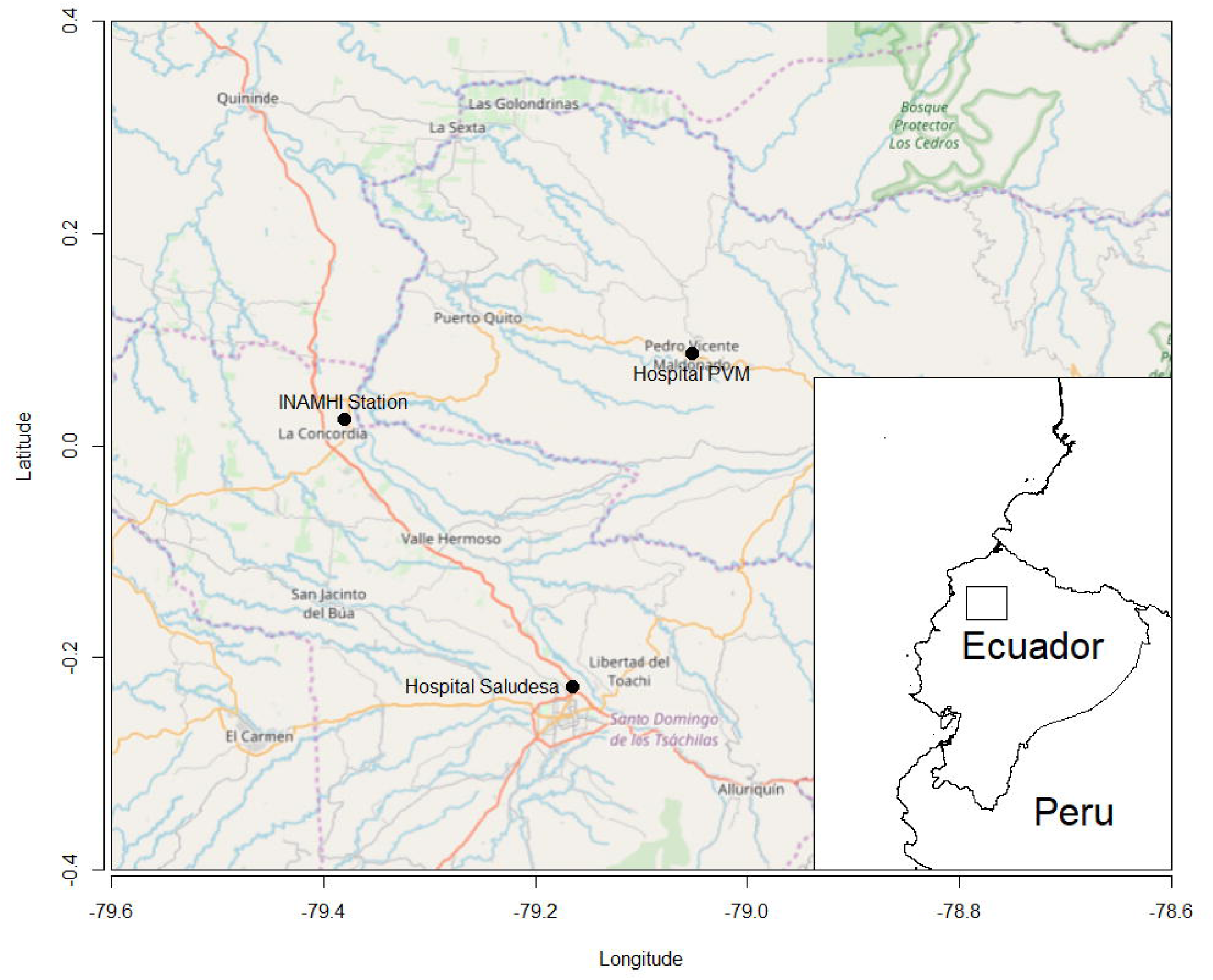
Study Site Locations. This map depicts the locations of the two hospitals used in the study, Hospital Saludesa and Hospital Pedro Vicente Maldonado, as well as the climate station. Inset, coast of Ecuador, with a square marking the relative position of the larger map. PVM=Pedro Vicente Maldonado, INAMHI=Instituto Nacional de Meteorología e Hidrología. Basemap tiles by © OpenStreetMap contributors, under CC BY-SA (https://www.openstreetmap.org/copyright). Inset tiles from by © Stamen Design, under CC BY 3.0 (maps.stamen.com). Maps were modified by R.S. for this manuscript.

### Data Collection

For this medical record review, we examined de-identified records with a primary diagnosis of arthropod-borne viral fevers and viral hemorrhagic fevers. These included International Statistical Classification of Diseases and Related Health Problems, 10th Revision (ICD-10) codes A90—A99. Records from Hospital Pedro Vicente Maldonado included consult dates from August 1, 2009 through July 31, 2016. Records from Hospital Saludesa included consult dates from July 1, 2014 through July 31, 2016. The following variables were available for analysis: consult date, primary diagnosis, ICD-10 code, and patient demographics (age, sex, insurance status, county-level address, weight, and height). We set criteria to exclude patients missing more than 50% of these variables. Information regarding school schedules and holiday dates in each year was obtained from the Ecuadorian Ministry of Education, the Ministry of Tourism and local residents [43-46]. School sessions and holidays analyzed in this study are in Table 1. Data for monthly climate variables measured at the La Concordia station (0°01′29.0″N, 79°22′49.0″W, Fig 1) were obtained from the National Institute of Meteorology and Hydrology in Ecuador [47, 48]. Oceanic Niño Indices (ONI), a measure of ENSO effects, were obtained from the National Weather Service (http://www.cpc.ncep.noaa.gov/products/analysis_monitoring/ensostuff/ensoyears.shtml).

**Table 1.**
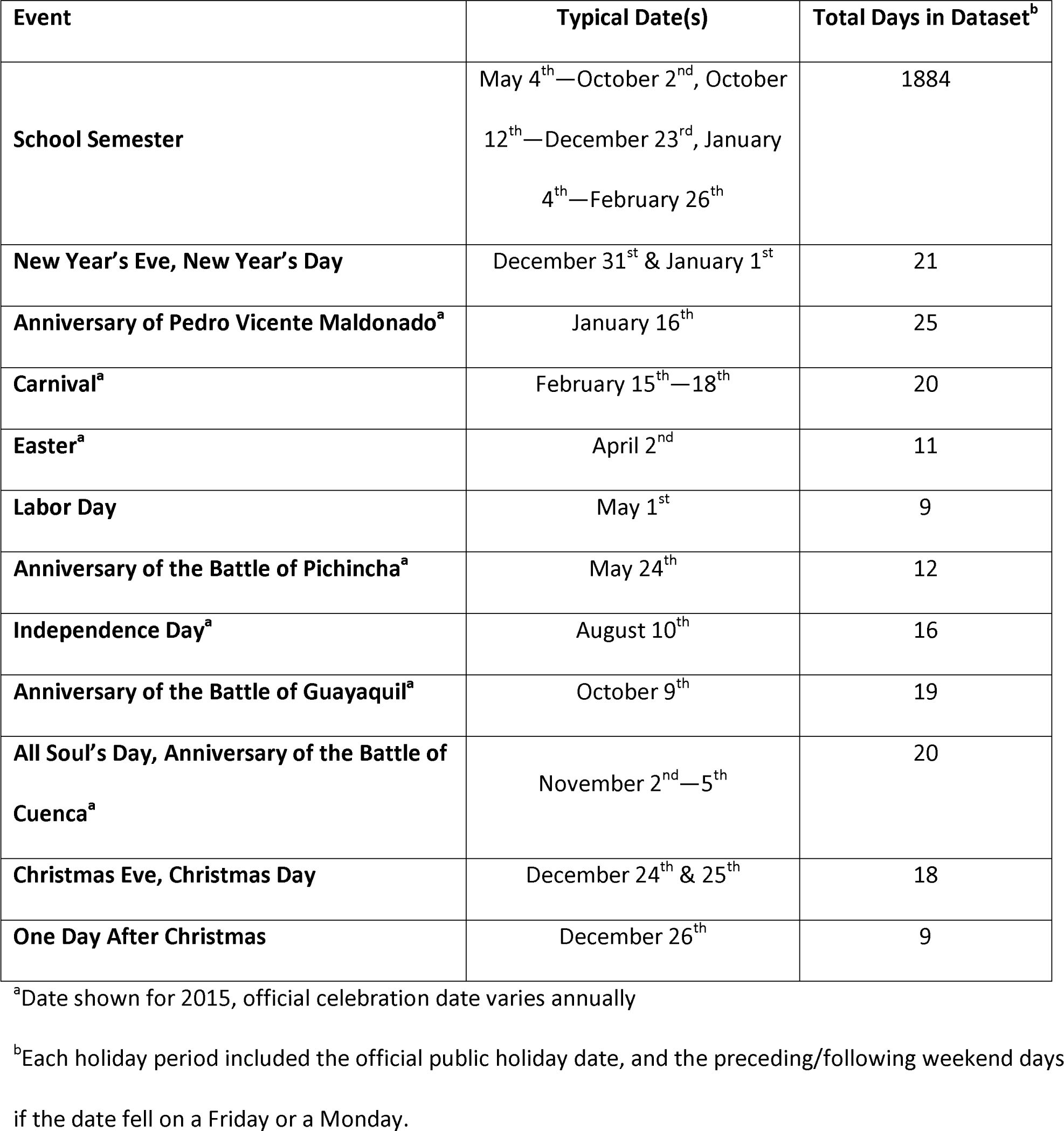
School Sessions & Holidays.

### Ethics

This research was certified as non-human subjects research by the Institutional Review Board of University of Wisconsin-Madison (#2017-0033).

### Statistical Analysis

Four observations (5% of total) for monthly absolute minimum temperature were missing. Multiple imputation was used to estimate these values using all available monthly climate variables, monthly case counts, and time. Ten imputations were performed with a fully conditional specification algorithm; parameters were pooled and used to obtain estimates.

Log-linked Poisson generalized linear models with generalized estimating equations (GEE) (autoregressive correlation structure) were used for all models. Models using GEE account for correlation in data, as is common in time series data. To account for temporal autocorrelation, cases were clustered by week of diagnosis. Model fit was assessed using quasi-likelihood under the independence model information criterion (QIC). Model 1 was used to evaluate the intra-annual and long-term seasonality of disease in a non-climate seasonal model; these seasonal components were included because there is evidence for intra-annual patterns in dengue diagnoses elsewhere in Ecuador [20] and our dataset had large year-to-year variations in diagnoses. Daily case counts were the outcome of interest and data from both hospitals were combined, with an indicator variable for hospital of origin. Long-term trends were estimated with a restricted cubic spline; number of knots was determined by best fit. For intra-annual effects, we compared sine and cosine waves with frequencies of once, twice, and/or three times in 365 days, using best fit to select the final fit. After selecting the best fit for the long-term and intra-annual effects, we added day-of-week and holidays as indicator variables, with the hypothesis that day-of-week may impact care-seeking decisions, and that patients may be less likely to seek care on a holiday (due to family obligations or travel). Holidays included official government-declared holidays and any weekends immediately before or after these holidays, as well as the day after Christmas.

Because children may differ in their exposure to dengue risk factors when school is in session, the effect of school schedules were examined using a subset analysis (Model 2). We restricted this analysis to school-aged children (ages 4—18) who sought care at Hospital Pedro Vicente Maldonado (n=142). We used the best-fit long-term and intra-annual effects model from the first analysis and included an indicator variable for days where school was in session (including weekends during the school year).

To determine the impact of climate on disease seasonality, we built a log-linked Poisson generalized linear model (Model 3). Climate data were available as monthly averages, so daily case counts were aggregated to monthly counts. Temperature and precipitation variables were centered on their mean value; temperatures were scaled at 2° Celsius, number of days with precipitation were scaled at 5 days and total monthly precipitation was scaled at 10 mm. The effects of climate variables (all continuous or integer variables), including ONI, average monthly temperature, minimum monthly temperature, maximum monthly temperature, total monthly precipitation, and number of days per month with precipitation were evaluated. Because climate variables interact with each other in reality, we also examined interactions between the significant climate variables in the final model.

Data analysis and visualization was performed using SAS version 9.2 (SAS Institute, Cary, NC) including the macros DASPLINE, DSHIDE, and weekno [49, 50], and R version 3.2.2 (R Foundation for Statistical Computing, Vienna, Austria) including packages haven, raster, dismo, ggmap, OpenStreetMap, sp, geepack, and MASS [51-59].

## Results

Characteristics of the data used in this study are in Table 2, with patient demographics available in Supplemental Table 1. No cases met the exclusion criteria; all cases were included in analysis. The diagnoses in the dataset included dengue fever (A90), dengue hemorrhagic fever (A91), other mosquito-borne viral fevers (A92), and mosquito-borne viral encephalitis (A83). Dengue diagnoses comprised 98.7% of the patients in the study. On average, one case is diagnosed at Pedro Vicente Maldonado every 4.3 days, and one case is diagnosed at Saludesa every 25 days. Time series plots of aggregated monthly case data from both hospitals, and monthly climate data are in Figure 2.

**Table 2.**
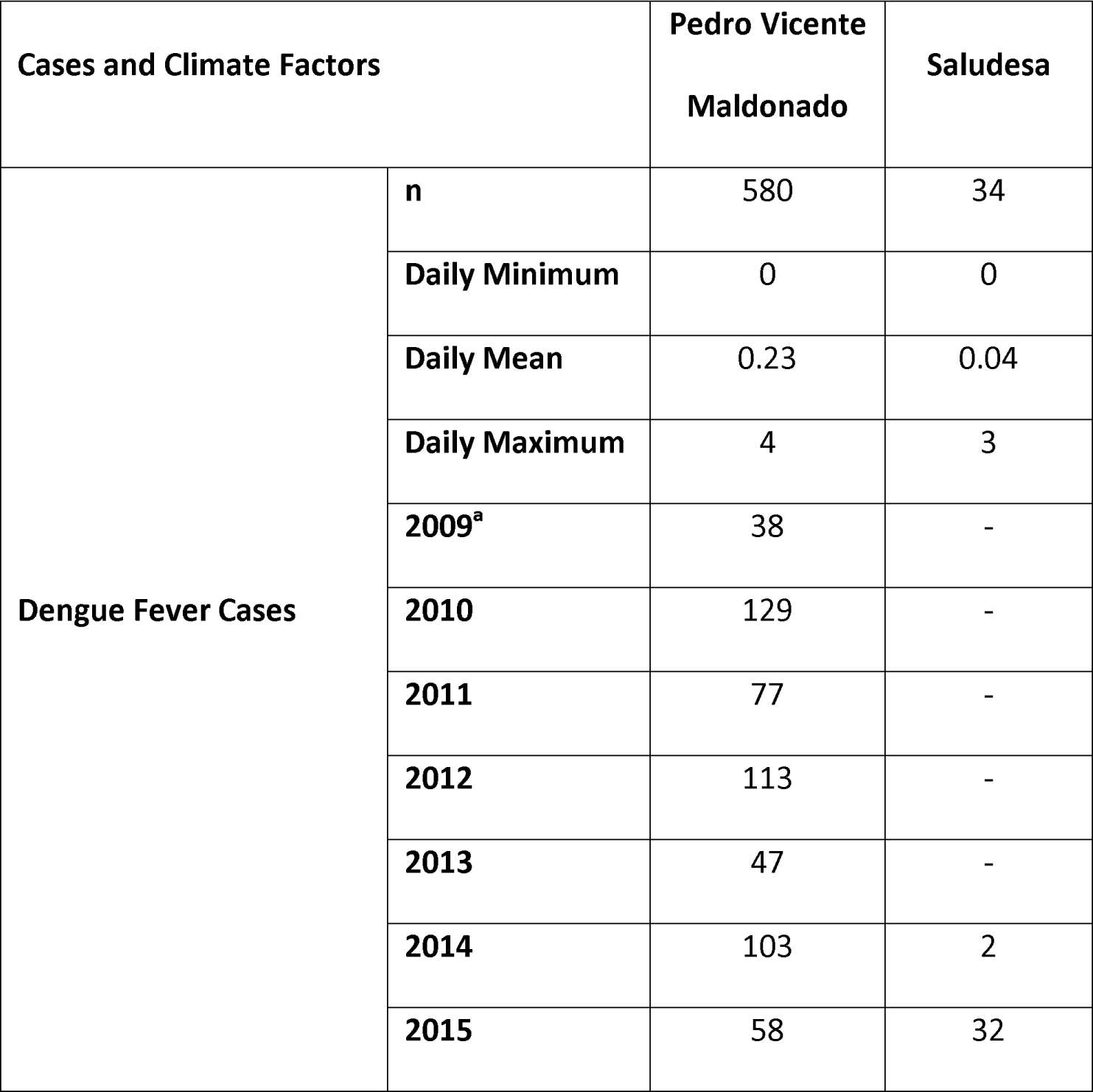
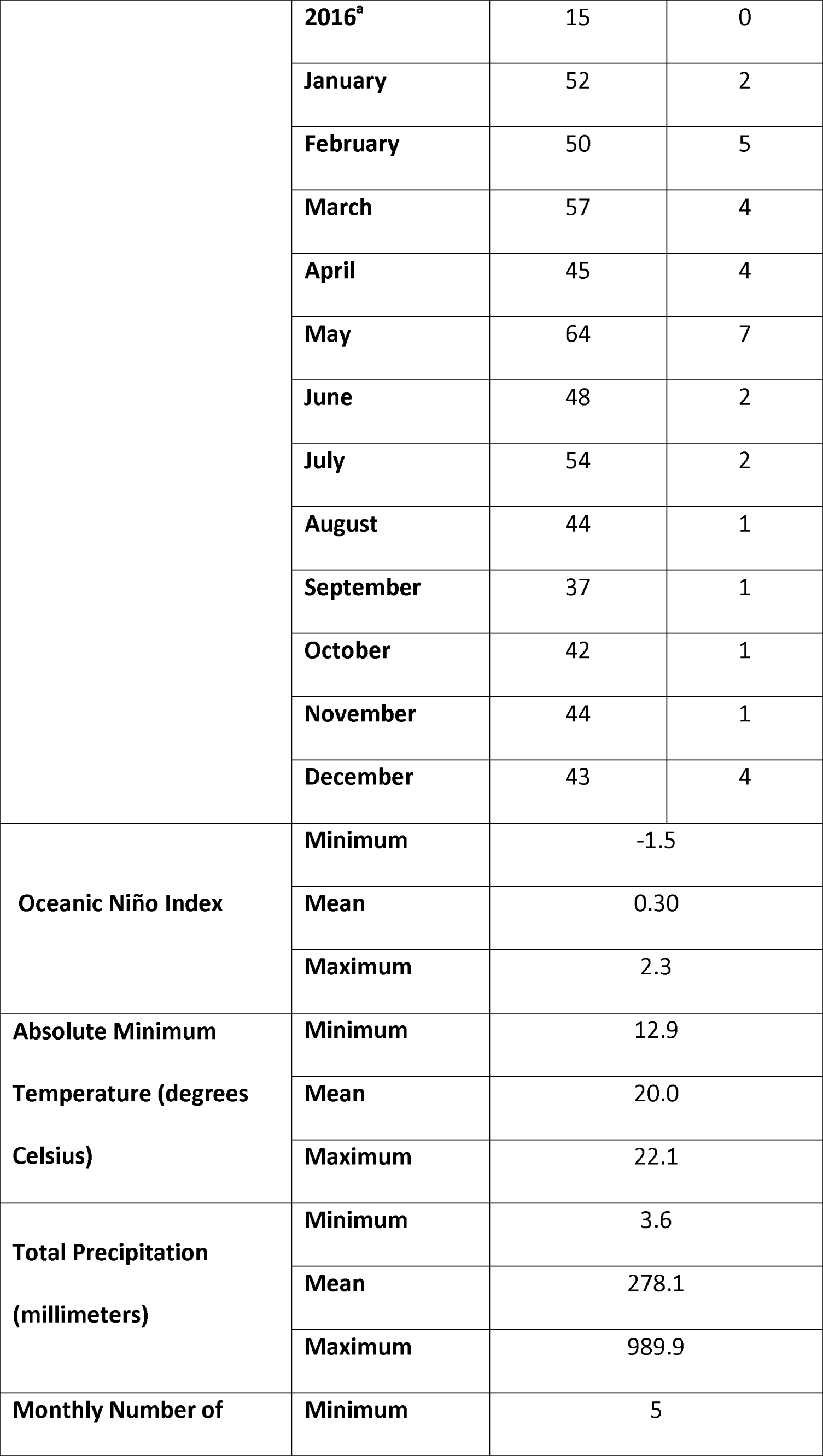
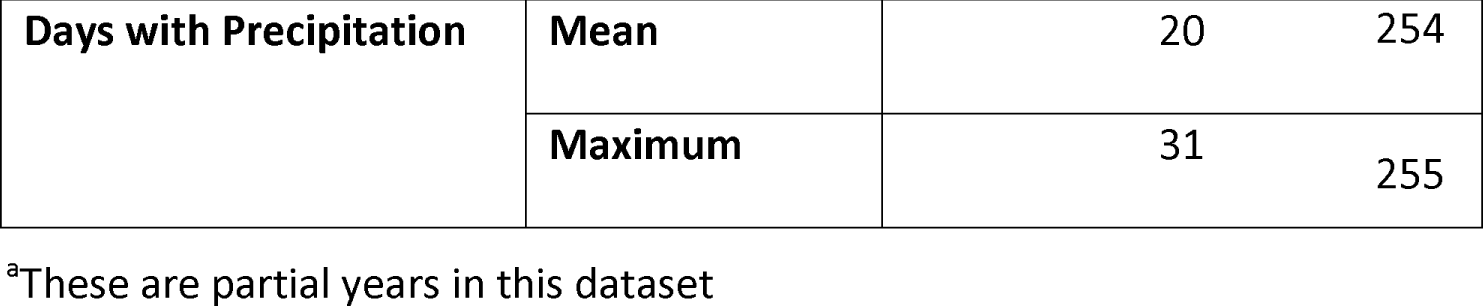
Data Characteristics.

**Fig 2.**
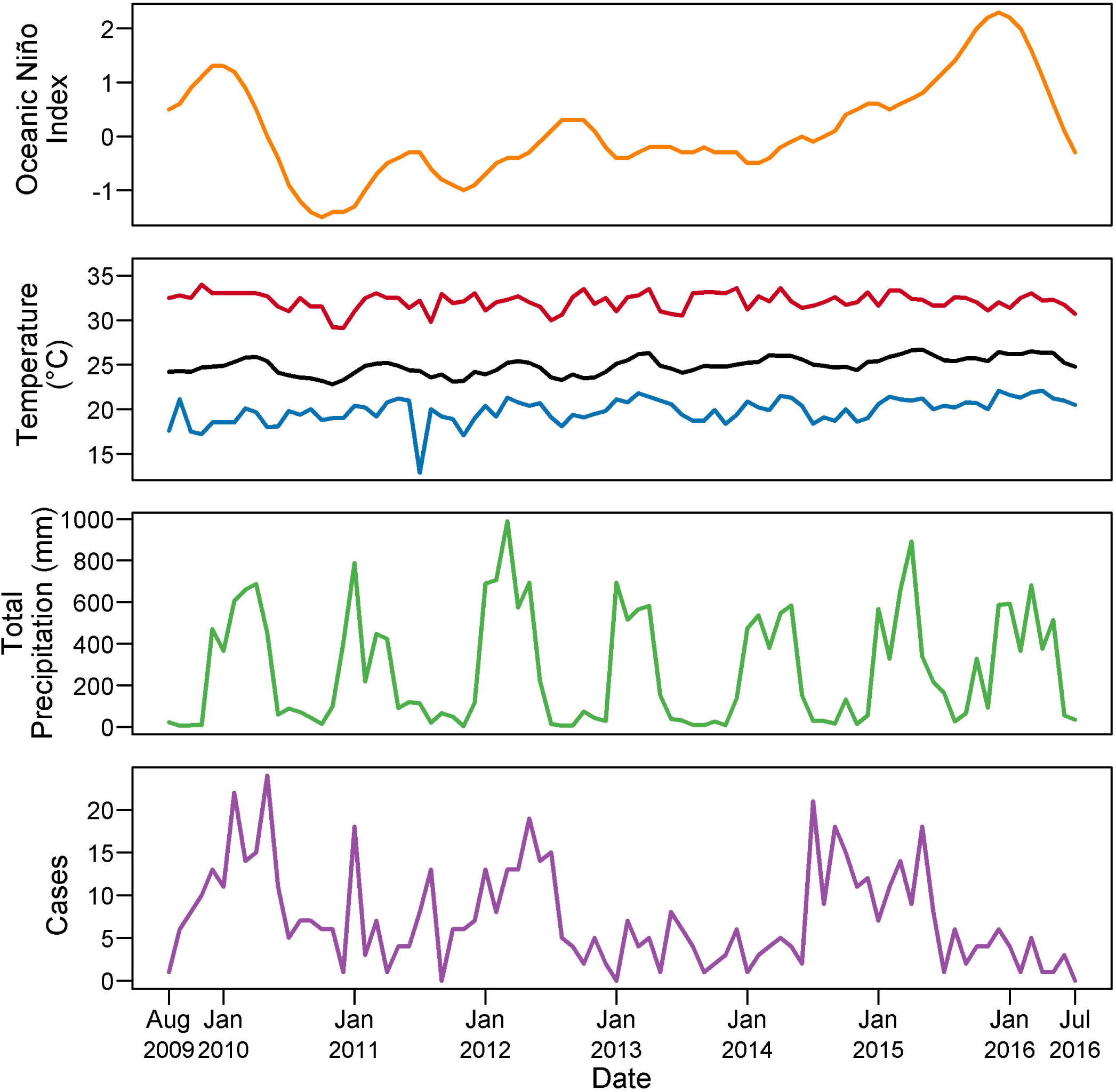
Time Series Plots for Dataset. Monthly averages for Oceanic Niño Index (orange), minimum (blue), mean (black), and maximum (red) temperature, precipitation (green), and diagnoses (purple) are plotted over the time period of the study.

The final model for non-climate seasonality (Model 1, parameters in Supplemental Table 2) included a sine and cosine wave with an annual cycle, long-term patterns, day-of-week effects, and indicator variables to designate holidays and hospitals. Fits metrics for the null model and each considered model are available in Supplemental Table 3. Model 1 predictions for daily diagnoses are presented in Fig 3 and exhibit an annual peak of disease in mid-March each year on average. Day-of-week effects are summarized in Fig 4. Compared to the average of all days, Tuesdays and Thursdays were more likely to have dengue fever diagnoses (Tuesday: relative risk (RR)=1.26, 95% confidence interval (CI) 1.05—1.51, p=0.012, Thursday: RR=1.25, 95% CI 1.02—1.52, p=0.033), while Saturdays and Sundays were less likely to have dengue fever diagnoses (Saturday: RR: 0.81, 95% CI 0.64—1.01, p=0.062 Sunday: RR: 0.74, 95% CI 0.58—0.95, p=0.016). Compared to non-holidays, dengue fever cases were much more likely to be diagnosed the day after Christmas (RR: 2.80, 95% CI 1.46—5.30, p=0.002), after holding all other covariates constant. The subanalysis (Model 2) did not find an effect of school session on dengue diagnoses.

**Fig 3.**
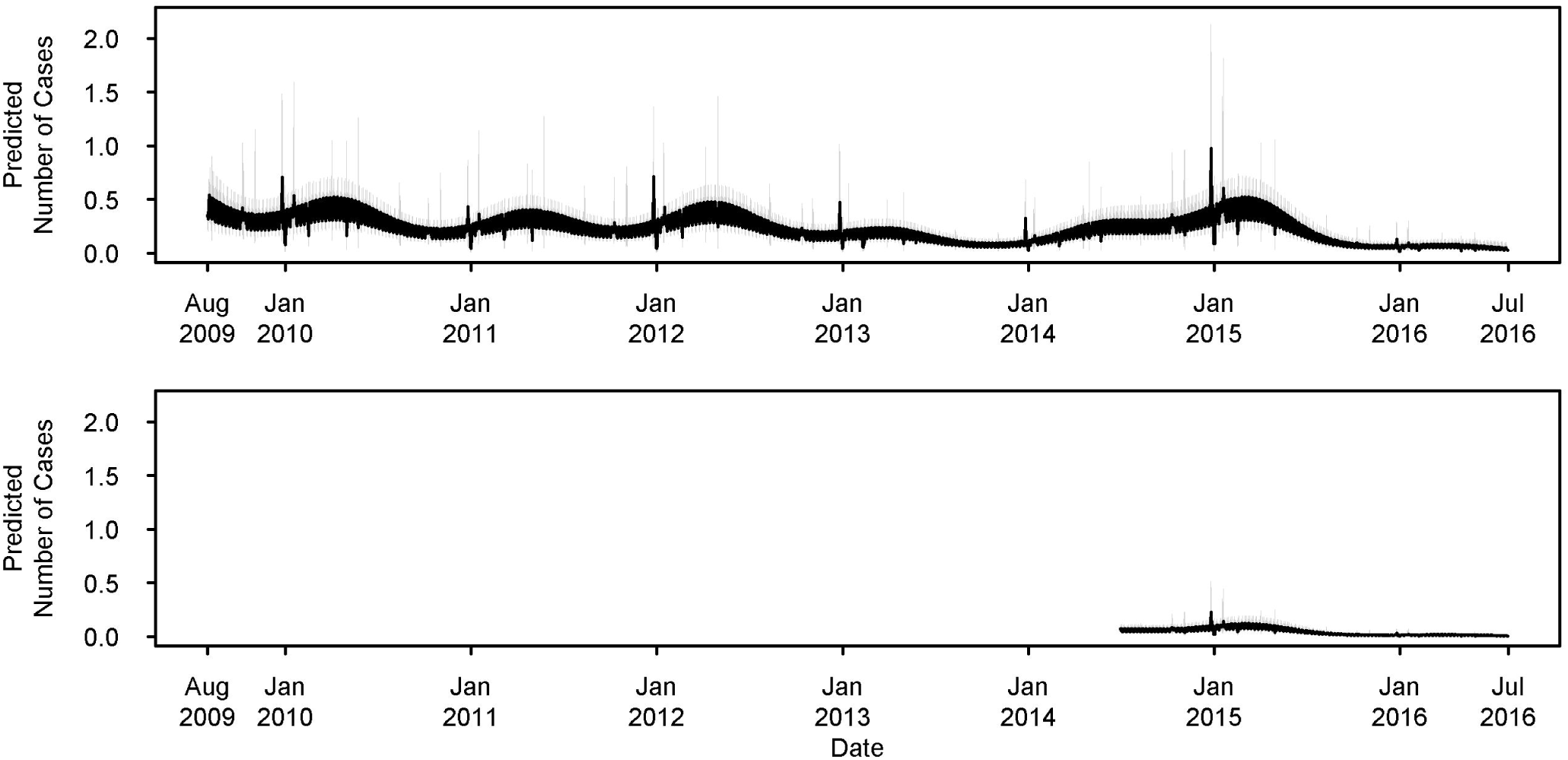
Non-climate Seasonality. The non-climate seasonality (Model 1) predictions for daily diagnoses exhibit an annual seasonality peaking in mid-March. Case predictions are in black, with confidence intervals in grey. The top panel depicts predictions for Hospital Pedro Vicente Maldonado and the bottom panel depicts predictions for Hospital Saludesa.

**Fig 4.**
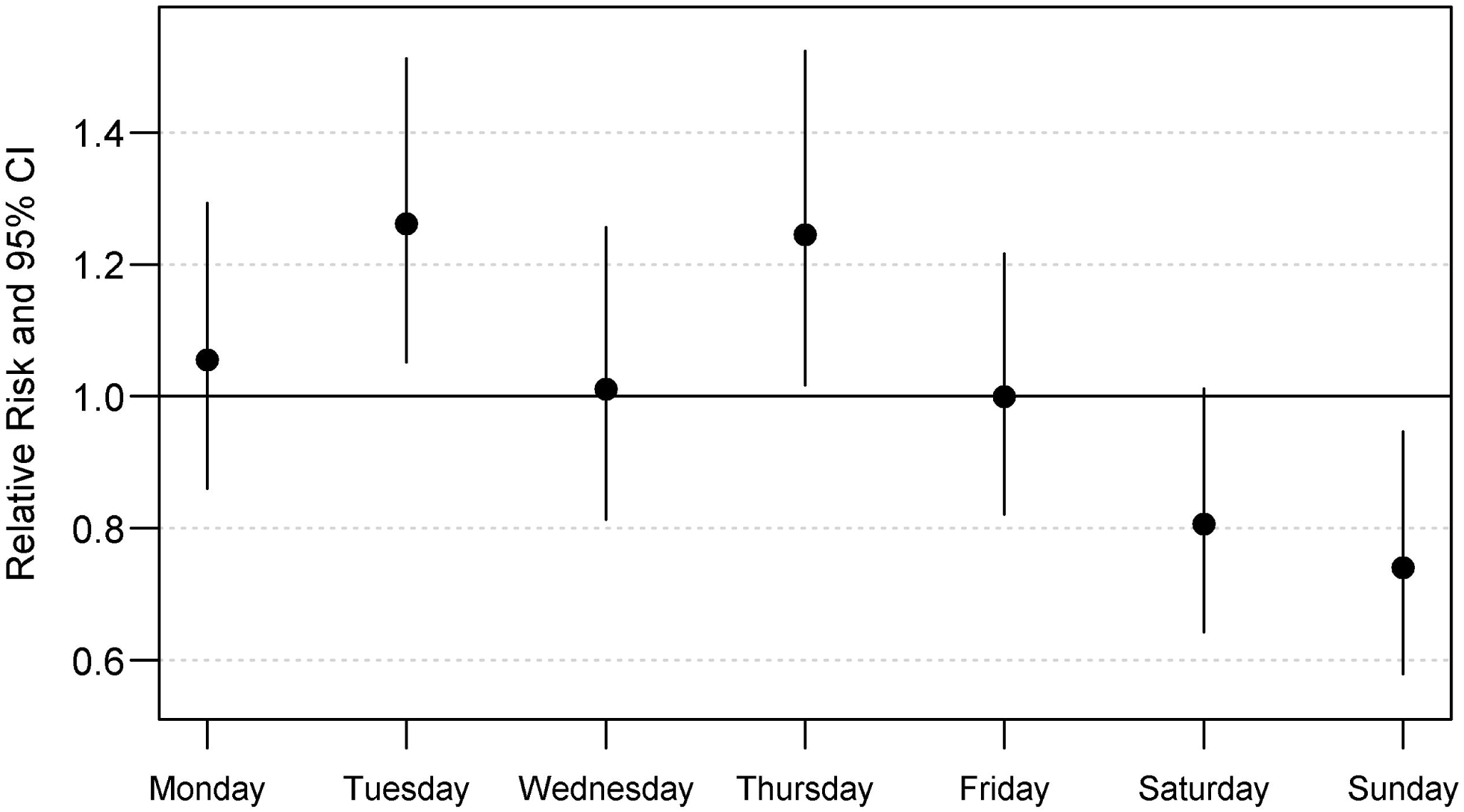
Day-of-Week Effects. The effect of the day of the week on dengue fever diagnoses is summarized in this graph, comparing each day to the overall average effect of weekday. A null estimate (RR=1.0) is included as a reference. Effect estimates are derived from Model 1. CI=confidence interval.

Most climate variables exhibited small but significant effects on risk of dengue fever diagnoses. Fit metrics for the null model and each considered climate model are available in Supplemental Table 4. Greater total monthly precipitation (RR: 2.14, 95% CI 1.26—3.64, p=0.005) results in increases in dengue fever diagnoses, *i.e*. for every 10mm increase in monthly precipitation, there is an approximately two-fold increase in dengue fever diagnoses on average. In addition, there were significant interactions between total monthly precipitation, number of days with precipitation, and monthly absolute minimum temperature. Model 3 predictions of interaction variable effects are in Fig 5, wherein observed values for monthly absolute minimum temperature, total monthly precipitation, and days per month with precipitation were used to predict the number of dengue cases per month within a reasonable range of precipitation and minimum temperature values. At an absolute minimum temperature of 18—19° C, the predicted number of cases increased (5 to 15 cases per month) as total monthly precipitation increased (from 125 to 875 mm per month) and *decreased* as the number of days with precipitation increased (from 5 to 30 days per month), but as minimum temperatures warm, the direction of these relationships changes. When the absolute minimum temperature is 20° C, additional days with precipitation or increases in monthly amounts of precipitation have little effect on the number of diagnoses. For a monthly minimum temperature of 21—22° C, the effect of increased amounts of precipitation is weaker, but still positive, while the impact of number of days with precipitation at warmer temperatures leads to *increases* in the number of dengue diagnoses (from 2 to 10 cases per month).

**Fig 5.**
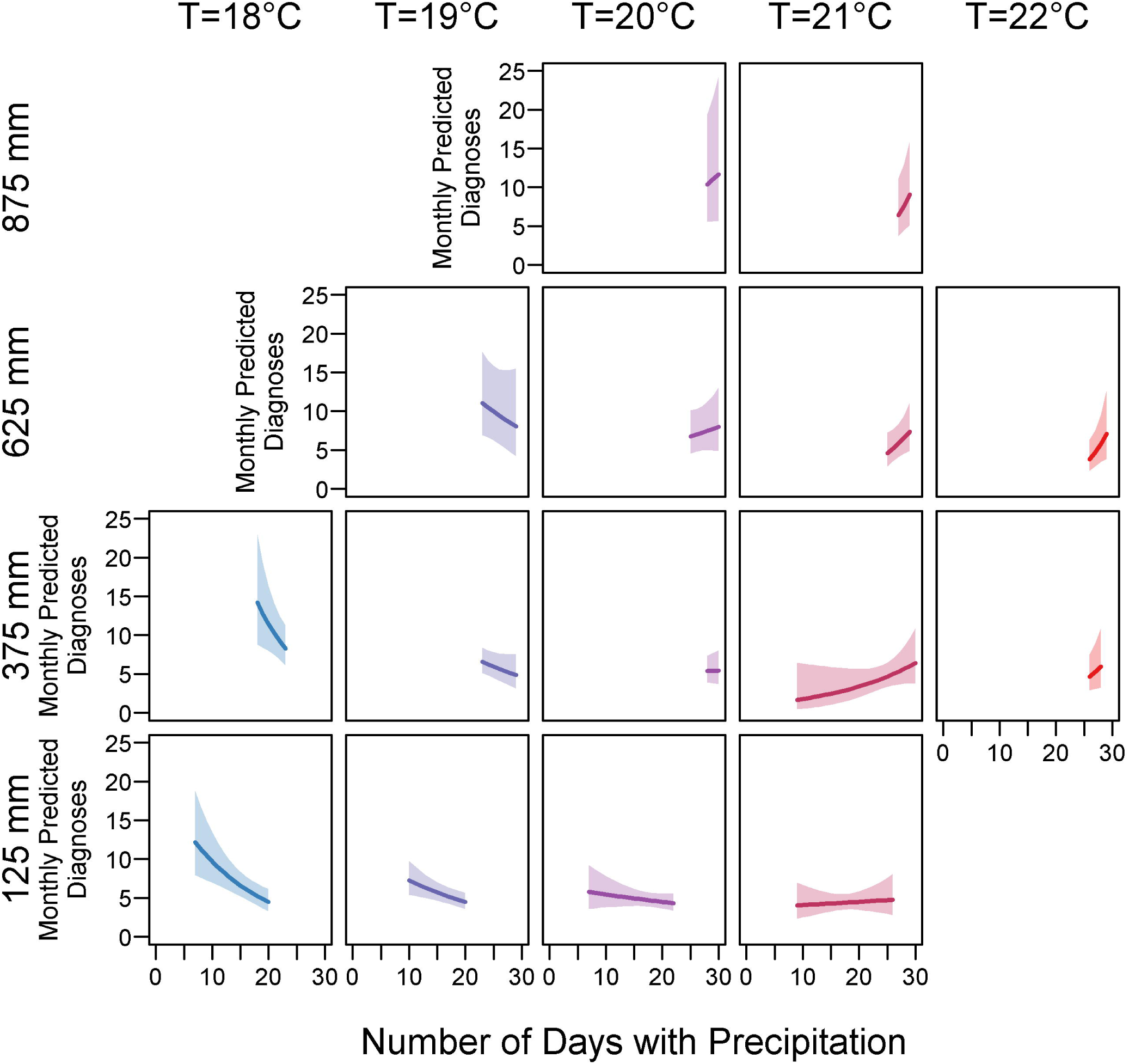
Interactions Between Monthly Precipitation and Monthly Minimum Temperature. Within individual plot panels, number of days with precipitation increase along the x-axis while monthly predicted number of dengue cases increase along the y-axis. Absolute minimum temperature levels increase along panel columns from left to right, and monthly amounts of precipitation increase along panel rows from bottom to top. Increases in the amount of precipitation leads to increases in the number of dengue diagnoses for all temperature conditions, but the relationship between temperature and number of days with precipitation exhibits an overall U-shaped pattern. As the minimum temperature warms, the relationship between number of days with precipitation and number of dengue diagnoses changes from negative to positive. At lower temperatures (18—19° C), additional days with precipitation lead to decreases in the predicted number of dengue cases. At 20° C, the relationship is flat, and at warmer temperatures (21—22° C), additional days with precipitation lead to increases in the number of dengue cases. Effect estimates were obtained from Model 3. T=monthly minimum temperature, mm=millimeters, C=Celsius.

## Discussion

Understanding the seasonality of infectious diseases can be crucial to the public health efforts to control these diseases. Seasonality is a major determinant of vaccination scheduling, timing of educational campaigns, and allocation of resources. In this paper, we examine non-climate (long-term trends, intra-annual patterns, day-of-week and holiday effects) and climate components of seasonality.

Our data exhibits annual peaks in dengue fever diagnoses, occurring in mid-March. Long-term studies of dengue seasonality in coastal Ecuador and other countries also exhibit annual peaks as well as inter-epidemic periods (high-intensity dengue fever seasons followed by a low-intensity season or seasons the following two years). Coastal Ecuador exhibits significant annual and two-year peaks in dengue incidence [20]; additional studies indicate that El Niño events, which occur in variable annual or multi-year patterns, may also influence dengue incidence patterns [19]. Our data exhibits peaks in 2012, and 2015; 2009 and 2015 were moderate and very strong were El Niño years, respectively [60]. Research from Peru suggests annual and three-year peaks in dengue incidence [61], while Colombia experiences two-to five-year cycles [62], with some parts of Colombia lacking annual disease peaks [63].

In this dataset, dengue fever diagnoses were likely affected by healthcare-seeking behavior. The decision and timing of seeking care for health problems can be affected by short-term time trends including day-of-week and holiday patterns. This type of research is scarce in South America. In the US and the UK, research on day-of-week effects has found that patients are less likely to visit the hospital on a weekend and that weekend hospital visits tend to be non-elective [64, 65], suggesting that patients may put off healthcare for less serious health conditions. Our findings agree with previous research, with Saturdays and Sundays being the least likely days for dengue fever diagnosis. However, we additionally found an increase in diagnoses on Tuesdays and Thursdays. We speculate that there may be some underlying pattern to diagnostic capabilities (e.g. staffing patterns, shipment days for lab supplies, or a backlog of patient samples from the weekend). We also examined holidays, with the reasoning that patients would also delay healthcare until after holidays. Previous studies suggest that holiday effects may be complex: research from Colombia has shown increases in dengue during periods immediately following holidays, from patients travelling to dengue-endemic areas during the holidays [66]. In our study, the individual holidays largely had no effect on dengue diagnoses, except for the day after Christmas (p=0.0015), when patients were more likely to be diagnosed with dengue fever. Since the incubation period for dengue is 4—10 days, and since most patients are local, we feel this spike in diagnoses is from those who became ill over the holiday and delayed their care, rather than acquired their illness during holiday travel. However, other than decreased diagnoses on Saturdays, day-to-day patterns of infectious disease healthcare-seeking at these hospitals did not have the same fluctuations as the dengue patients (data not shown), meaning these patterns may be the result of statistical noise and not general health-care seeking behaviors in the community.

In our seasonality assessment, we found that dengue fever diagnoses peaks during mid-March on average. This is the first assessment of dengue fever seasonality in rural Ecuador. Reports from nearby Colombia regarding dengue fever seasonality have not found an annual seasonal pattern for dengue incidence [63, 67], though these studies did not utilize sinusoidal variables, making it difficult to detect these patterns.

Climatic factors such as temperature or precipitation can affect the survival and distribution of mosquito vectors and the transmissibility of pathogens from these vectors [16-18]. In previous research in Colombia, studies have found average temperature, changes in average temperature, average relative humidity, total precipitation, and El Niño events to be major predictors of dengue incidence [63, 68]. Research in Ecuador has been limited to studies of dengue cases in coastal regions. In one study, minimum weekly temperature and weekly average precipitation were shown to be strongly linked to weekly number of dengue cases [20]. Minimum weekly temperature, precipitation, and El Niño events were also positively associated with dengue risk [19]. Our data illustrate a complex relationship between climate factors and dengue fever diagnoses. Temperature is a major factor; dengue transmission is sensitive to extremes of temperature as *Aedes aegypti* propagate and transmit dengue best between 18—32° C [63], but precipitation is also important. In isolation, total monthly precipitation and number of days with precipitation had opposite effects, suggesting that sufficient precipitation is necessary for dengue cases to occur, but that too many days with precipitation decrease risk. However, when we consider minimum monthly temperature, temperature modifies the effects of precipitation in a U-shaped pattern. All amounts of precipitation drive increases in dengue diagnoses but additional days with precipitation lead to decreases in dengue diagnoses while when temperatures are lowest, as in the months of July through November (mean minimum temperatures of 19.0—19.7° C). During these months precipitation amounts are all below 250 mm on average and durations are 11.7 to 15.7 days on average, resulting in a relatively low predicted number of dengue fever cases. At warmer temperatures, both number of days with and amount of precipitation have positive relationships with the number of dengue diagnoses. At the warmer part of the year – *i.e*. December through June (mean minimum temperatures of 19.8—21.2° C), precipitation quantity is higher (mean 271.6—635.4 mm per month) and occurs on more days (mean 21.6—28.3 days per month).

Our results likely reflect the effect of precipitation on mosquitoes: female *Aedes aegypti* mosquitoes tend to lay eggs just above the water surface in containers [69] until additional precipitation (*i.e*. flooding of the eggs) causes the eggs to hatch, but too much precipitation can wash eggs or larvae out of their containers [70], meaning some dry periods are necessary or even beneficial to *Aedes aegypti* abundance. Previous research has found that *Aedes aegypti* breeding site occupancy is increased at sites with longer dry periods [71]. Temperature levels affect evaporation rates and the durability of standing water (*i.e*. breeding and development sites); this may explain temperature’s modifying impact on the relationship between precipitation and dengue diagnoses.

Human hosts may also change their travel outside the home during consistently rainy periods, which may alter their exposure to dengue-infected mosquitoes (depending on where they are most exposed). Research in Australia found that virus acquisition was spatiotemporally linked to the case’s residence in 42% of dengue cases [72], though this proportion may differ in other geographic locations. Human movement and behavior is a major component of dengue fever risk [73]. Weather patterns affect human movements, with high movement variation on days with higher precipitation [74]. The patterns between dengue fever risk and climate variables observed in our data are likely a combination of the effect of climate on mosquito vectors and human behaviors.

Notably, the best-fit climate seasonality model included both long-term and annual sinusoidal variables, in addition to climate variables. If dengue seasonality were entirely driven by climate, we would expect that a model adjusting for the effects of climate to be sufficient with no long-term nor annual sinusoidal variables (*i.e*. all variation in the diagnosis rate would be explained by the climate variables). The importance of the long-term and annual sinusoidal variables in our climate model suggests that we are not completely adjusting for the effect of climate or that non-climate phenomena impact the seasonality of dengue diagnoses. Our ability to disentangle climate and non-climate seasonality is complicated by the introduction of chikungunya into a naïve population in 2015. These cases were treated as dengue diagnoses in our models, but it is impossible to know how many during this period were true dengue cases. Because chikungunya and dengue are spread by the same mosquito species, *Aedes aegypti*, we expect that many of the effects of climate will be the same for both chikungunya and dengue. The effects of chikungunya emergence on overall seasonality are important to consider. This introduction occurred outside of the typical dengue season (November) and had a high number of cases, meaning that the average annual peak of dengue is slightly earlier in our model than the true average annual peak of dengue in this population.

### Limitations

This dataset combines the patients from two hospitals. The patients at each hospital differ in their gender composition and insurance status. Both gender and insurance status likely affect healthcare-seeking behavior, meaning that the hospital populations may have different non-climate seasonal patterns of dengue diagnoses and some uncontrolled confounding may affect our results. However, we do control for the source hospital in our analysis which would control for some of these differences, and since the majority of cases (94.5%) are from one hospital, we do not expect this issue to substantially affect our results.

This dataset represents dengue fever diagnoses in the community and is only a proxy for dengue fever incidence rates. There are likely to be many more cases dengue fever in the community: 80% of dengue cases are estimated to be asymptomatic, some symptomatic patients may never seek care, and some symptomatic patients may have sought care at hospitals other than those included in this study. This could be a potential source of selection bias. However, our study hospitals are the major source of care in their communities and we are assessing seasonality and climate variables; we have no reason to believe that the effect of seasonality and climate is any different among symptomatic versus asymptomatic patients nor for the small number of persons who sought care at other clinics. The effect of selection bias on these data is likely minimal.

Dengue diagnosis can be difficult even for experienced clinicians, especially in a resource-limited setting such as Ecuador. Not all patients with a final dengue diagnosis were necessarily lab-confirmed; the use a laboratory confirmation likely varies by clinician, patient, and presenting symptoms, though the clinicians at the study hospitals are all experienced with dengue diagnosis. Based on observed hospital practices, we believe many of the cases in our dataset had positive dengue rapid tests, but that some were clinically diagnosed. Because not all cases were laboratory-confirmed, it is possible that some non-dengue cases were diagnosed as dengue, particularly when chikungunya was introduced to Ecuador (late 2015) and no diagnostic tools were available for chikungunya.

Our dataset only covers a seven-year period making it difficult to conclude if our observations truly reflect long-term or multi-year disease trends in this community. Additional research for longer periods of time will reveal if three-year peaks or changes in dengue fever diagnose rates are present in this community. The diagnosis of dengue could have been impacted by additional phenomena over the study period. Changes in mosquito control practices could affect actual disease rates or worsening economic conditions in Ecuador (due to a decrease in oil prices) would adversely affect the ability of patients to seek healthcare. In addition, a major earthquake in April 2016 disrupted many services in Ecuador, including transportation, utilities, and healthcare for several weeks, which may have disrupted the typical healthcare-seeking behavior of patients and the diagnostic capabilities of the hospitals during this time.

Available climate data was captured from a climatological station located 39 and 36 kilometers from Hospital Pedro Vicente and Hospital Saludesa, respectively. These data are only a proxy for actual climate conditions in our communities of interest. In addition, analyses with climate variables were limited to monthly summaries of these variables, making it difficult to ascertain if the relationships discovered in this research reflect the true relationship between climate variables and dengue fever diagnoses in these populations. Under the assumption that most patients would be bitten, experience symptoms, and seek care within the same month, the climate-diagnosis relationships presented in this study are a good estimate of dengue seasonality in these communities. In reality, there is considerable variation among the climate variables, mosquito exposure and dengue diagnoses in this community, which we were unable to capture in this study. Nor are we able to estimate the effects of climate variable interactions among ranges and combinations of variables that were unobserved in this location. In addition, the effect estimates for the climate variable interactions were often based on a small sample size, leading to wide confidence intervals for these estimates. Indeed, this veracity of this interaction will need to be confirmed with additional research. Future research will also address the limited range and unobserved climate combinations in this dataset by testing this interaction with data from areas with different climate conditions.

## Acknowledgements

Thank you to the information technology staff at Hospital Pedro Vicente Maldonado and Dr. Daniel Larco for explanation of diagnostic methods at the sites.

---

**Supplemental Figure:**
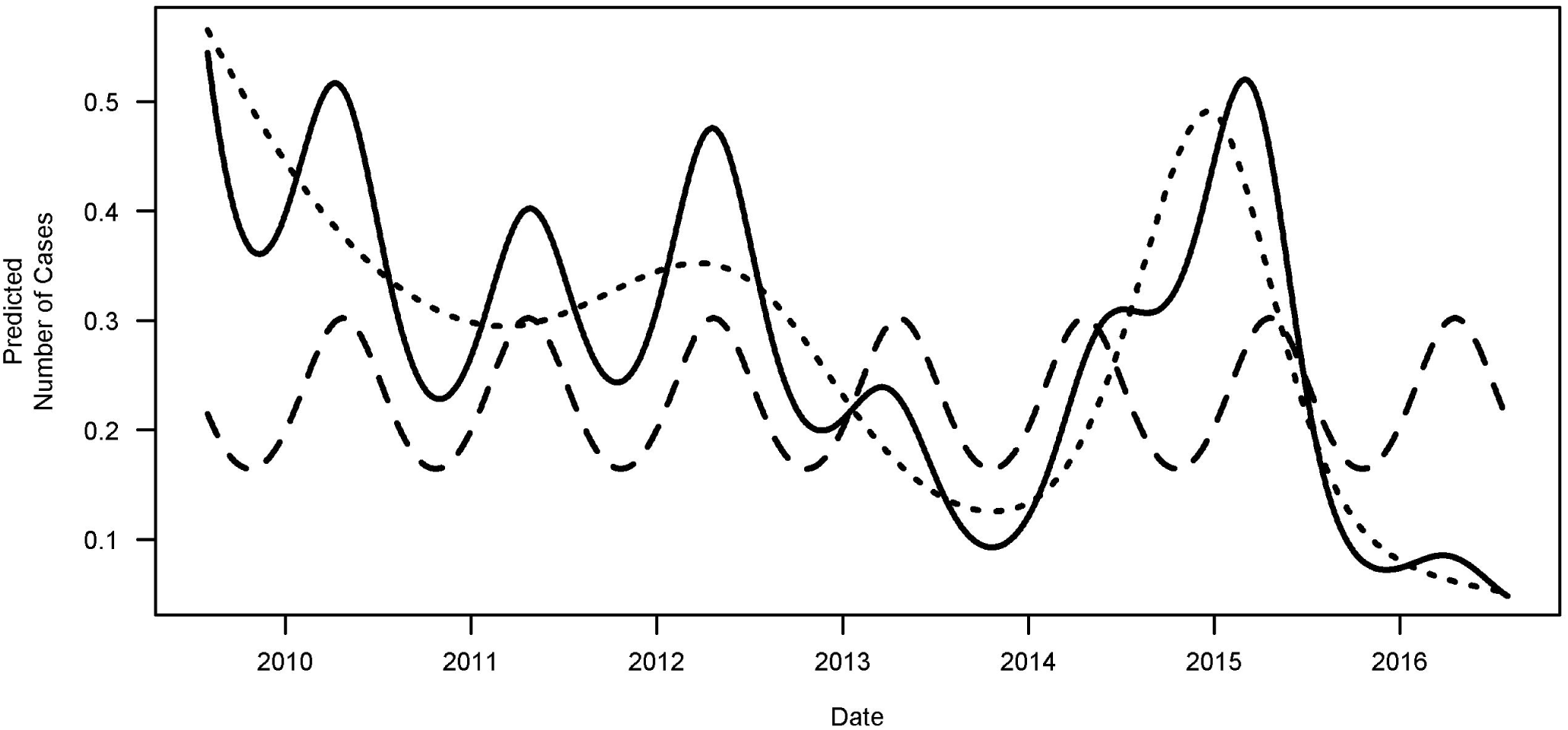
Splines and Sine/Cosine Effects in Model 1. The dashed line is the combined effect of the sine and cosine effects in the model, representing the annual fluctuation of dengue. The dotted line is the effect of the 7-knot spline, representing the long-term or inter-annual fluctuation of dengue. The solid line is the combination of these two effects in Model 1.

