## Supplemental Table 1 for "Seasonal patterns of dengue fever in rural Ecuador: 2009—2016 Seasonality of dengue fever in Ecuador"

**Supplementary Table 1: Patient Characteristics.**

Characteristics of the patient populations in this study are summarized, with hospital-specific percentages representing the proportion of each hospital’s population with each characteristic, the total percentage representing the proportion of the total (both hospitals) with each characteristic.

| Characteristics | | HPVM^a^ | Saludesa | Total |
| --- | --- | --- | --- | --- |
|  |  | Count  (%) | Count  (%) | Count  (%) |
| n | | 580  (94.5) | 34  (5.5) | 614  (100) |
| Gender | Male | 355  (61.2) | 9  (26.5) | 364  (59.3) |
|  | Female | 225  (38.8) | 25  (73.5) | 250  (40.7) |
| Age Groups (years) | 0—9 | 83  (14.3) | 3  (8.8) | 86  (14.0) |
|  | 10—19 | 106  (17.3) | 10  (29.4) | 116  (18.9) |
|  | 20—29 | 141  (24.3) | 8  (23.5) | 149  (24.3) |
|  | 30—39 | 121  (20.9) | 2  (5.9) | 123  (20.0) |
|  | 40—49 | 64  (11.0) | 6  (17.7) | 70  (11.4) |
|  | 50—59 | 38  (6.6) | 2  (5.9) | 40  (6.5) |
|  | 60—69 | 15  (2.6) | 1  (2.9) | 16  (2.6) |
|  | 70+ | 12  (2.1) | 2  (5.9) | 14  (2.3) |
| Diagnosis | Dengue fever | 570  (98.3) | 30  (88.2) | 600  (97.7) |
|  | Dengue hemorrhagic fever | 5  (0.9) | 1  (2.9) | 6  (1.0) |
|  | Mosquito-borne viral encephalitis | 1  (0.2) | 1  (2.9) | 2  (0.3) |
|  | Other mosquito-borne viral fever | 4  (0.7) | 2  (5.9) | 6  (1.0) |
| Address | Within same county as source hospital | 326  (56.2) | 28  (82.4) | 354  (57.7) |
|  | Within county adjacent to source hospital | 212  (36.6) | 3  (8.8) | 215  (35.0) |
|  | County no adjacent to source hospital | 31  (5.3) | 2  (5.9) | 33  (5.4) |
|  | Unknown | 11  (1.9) | 1  (2.9) | 12  (2.0) |
| Insurance Status | Private | 1  (0.2) | 1  (2.9) | 2  (0.3) |
|  | IESS | 477  (82.2) | 0  (0.0) | 477  (77.7) |
|  | None | 100  (17.2) | 28  (82.4) | 128  (20.8) |
|  | Other | 1. 2 2. (0.3) | 5  (14.7) | 7  (1.1) |

^a^HPVM=Hospital Pedro Vicente Maldonado
