## Supplemental Table 2 for "Seasonal patterns of dengue fever in rural Ecuador: 2009—2016 Seasonality of dengue fever in Ecuador"

**Supplementary Table 2.**

Effect estimates and 95% confidence intervals for Model 1. Both effect estimates and 95% confidence intervals have been converted to rate ratios for ease of interpretability.

| **Parameter** | **Estimate** | **95% Confidence Interval** | | **p-value** |
| --- | --- | --- | --- | --- |
| **Intercept** | 0.08 | 0.04 | 0.15 | <.0001 |
| **sin(2πt)** | 0.74 | 0.65 | 0.85 | <.0001 |
| **cos(2πt)** | 0.97 | 0.84 | 1.12 | 0.6738 |
| **k^1^** | 1.00 | 1.00 | 1.00 | 0.0057 |
| **k1** | 1.01 | 1.00 | 1.02 | 0.0068 |
| **k2** | 0.95 | 0.92 | 0.98 | 0.0006 |
| **k3** | 1.14 | 1.08 | 1.20 | <.0001 |
| **k4** | 0.69 | 0.61 | 0.79 | <.0001 |
| **k5** | 1.85 | 1.39 | 2.46 | <.0001 |
| **Monday^2^** | 1.05 | 0.86 | 1.29 | 0.6068 |
| **Tuesday** | 1.26 | 1.05 | 1.51 | 0.0122 |
| **Wednesday** | 1.01 | 0.81 | 1.26 | 0.9204 |
| **Thursday** | 1.25 | 1.02 | 1.52 | 0.0333 |
| **Friday** | 1.00 | 0.82 | 1.22 | 0.9965 |
| **Saturday** | 0.81 | 0.64 | 1.01 | 0.0629 |
| **Sunday** | 0.74 | 0.58 | 0.95 | 0.0164 |
| **Hospital PVM** | 4.28 | 2.67 | 6.88 | <.0001 |
| **NYE** | 0.26 | 0.04 | 1.60 | 0.145 |
| **An** | 1.50 | 0.50 | 4.51 | 0.4729 |
| **Carnival** | 0.42 | 0.13 | 1.37 | 0.1519 |
| **Easter** | 0.83 | 0.26 | 2.60 | 0.7488 |
| **Labor Day** | 0.51 | 0.08 | 3.06 | 0.4594 |
| **Pichincha** | 0.60 | 0.11 | 3.27 | 0.5521 |
| **Independence** | 1.05 | 0.40 | 2.73 | 0.9213 |
| **Guayaquil** | 1.38 | 0.59 | 3.29 | 0.4598 |
| **All Souls** | 0.97 | 0.30 | 3.18 | 0.9634 |
| **Christmas** | 1.29 | 0.35 | 4.74 | 0.6985 |
| **Day after Christmas** | 2.80 | 1.48 | 5.30 | 0.0015 |

^1^Parameters k through k5 are the spline parameters representing long-term trends in the model.

^2^The effects for days of the week were calculated in comparison to the average effect of all days.

Hospital PVM=Hospital Pedro Vicente Maldonado, NYE=New Year’s Eve, An=Anniversary of the Founding of Pedro Vicente Maldonado
