## Supplemental Table 3 for "Seasonal patterns of dengue fever in rural Ecuador: 2009—2016 Seasonality of dengue fever in Ecuador"

| **Model** | **Variables** | **QIC^a^** |
| --- | --- | --- |
| Null model | intercept only | 2606 |
| Hospital | Indicator for hospital | 2559 |
| Long-term | 7-knot spline, hospital | 2540 |
| Long-term & annual short-term | 7-knot spline, sin and cosine (f=365 days), hospital | 2509 |
| Long-term & biannual short-term | 7-knot spline, sin and cosine (f=182.5 days), hospital | 2560 |
| Long-term & three times annual short-term | 7-knot spline, sin and cosine (f=91.25 days), hospital | 2527 |
| Final model | 7-knot spline, sin and cosine (f=365 days), hospital, weekdays and holidays | 2533 |

**Supplementary Table 3.**

Descriptions and metrics for models considered during the analysis of temporal seasonality in the dataset (Model 1).

^a^QIC=quasi-likelihood under the independence model information criterion
