## Supplemental Table 4 for "Seasonal patterns of dengue fever in rural Ecuador: 2009—2016 Seasonality of dengue fever in Ecuador"

| **Model** | **Variables** | **QIC^a^** |
| --- | --- | --- |
| Hospital & temporal | 7-knot spline, sin and cosine (f=365 days), hospital | -1167 |
| Climate variables | 7-knot spline, sin and cosine (f=365 days), hospital, climate variables | -1186 |
| Final model | 7-knot spline, sin and cosine (f=365 days), hospital, climate variables and interactions | -1203 |

**Supplementary Table 4.**

Descriptions and metrics for models considered during the analysis of climate seasonality in the dataset.

^a^QIC=quasi-likelihood under the independence model information criterion
